# Frontoparietal effective connectivity supports working memory in early childhood

**DOI:** 10.64898/2026.09.23.753700

**Authors:** Carole Guedj, Lia Bollinger, Damien Marie, Gianvito Laera, Romain Ceresetti, Alessio Comparini, Jonas Richiardi, Matthias Kliegel, Clara E. James

## Abstract

Working memory (WM), a core executive function that supports learning and reasoning, undergoes rapid development during early childhood years. Neural activation patterns underlying WM have been extensively studied in both adults and children, revealing increasing involvement of the frontoparietal network (FPN) with age and coordinated interactions with the default mode network (DMN). However, relatively little is known about how these network interactions develop in children, particularly within the visuospatial domain. The present study investigated the influence of age and task performance on effective connectivity within the FPN and its interactions with DMN hubs during a visuospatial WM task in early childhood. Functional MRI data were collected during a visual n-back task with two WM load levels (0-back and 2-back) in 120 typically developing children aged 5-8 years. Dynamic Causal Modeling (DCM) with parametric empirical Bayes was used to estimate effective connectivity among eight regions of interest spanning the FPN and DMN. Behaviorally, age-related gains were steepest under high-load for accuracy, and under low-load for reaction time, suggesting WM capacity is a demand-sensitive marker of development at this age. At the network level, high-load robustly engaged the FPN, whereas low-load was associated with relatively greater engagement of DMN and episodic memory-related regions. Baseline effective connectivity was dominated by parietal-to-frontal drive and strong self-inhibition in the anterior cingulate cortex. Age-related effects on this architecture were sparse, involving asymmetric interhemispheric frontal coupling and reduced parieto-cingulate coupling, whereas accuracy-related effects were larger in magnitude, particularly at self-connections. More accurate children showed greater excitability in frontal and parietal FPN nodes alongside increased self-inhibition in the precuneus, a DMN hub, consistent with a neural efficiency account in which better performance reflects more selective, rather than more global, network engagement. Task load further shaped connectivity in distinct ways: low-load elicited widespread inter-regional modulation, whereas high-load predominantly modulated self-connections, with age and accuracy effects on this modulation dissociating by load. These findings suggest that, in primary school children, current task performance was associated with more pronounced differences in frontoparietal effective connectivity than was age, and that WM-related network engagement becomes increasingly selective and load-dependent across this developmental window.

## Introduction

Working memory (WM) is a core executive function (EF) referring to the ability to temporarily maintain, manipulate and update goal-relevant information (Baddeley, 1992; Miyake et al., 2000). It is involved in higher-order cognitive processes such as planning, reasoning, problem-solving and self-regulation (Diamond, 2013). WM also supports language acquisition (Fiske & Holmboe, 2019) as well as arithmetic skills and broader academic achievement (Gathercole et al., 2004; Rosenberg et al., 2020). Accordingly, many studies consider the development of WM as a key driver of overall cognitive growth (Fiske & Holmboe, 2019; Hartung et al., 2020; Kail, 1985). WM improves steeply during early childhood years, paralleling rapid brain growth and network reorganization, and continues to mature into adolescence (Diamond, 2002, 2013). Understanding this developmental window is critical both for characterizing typical cognitive trajectories and for early identification of difficulties. Intervention studies targeting executive functions such as WM directly (e.g. working memory training, Sala & Gobet, 2020), or indirectly (e.g. music interventions, (James et al., 2020) rest on mechanistic and developmental insights (Diamond & Lee, 2011). Yet how WM develops at the onset of formal schooling and the relationship with brain maturation remain to be further established.

Within WM, a useful distinction can be drawn between verbal and visuospatial domains. Baddeley and Hitch’s multicomponent model (1974) divides WM into four components: a central executive (CE) that monitors two modality-specific subsystems: the phonological loop (PL, for phonological/verbal information) and the visuospatial sketchpad (VSSP, for visuospatial information), and an episodic buffer, which links information from these subsystems to long-term memory. Most studies treat verbal/auditory and visuospatial WM as separate subsystems, a distinction that has helped identify partially divergent neural substrates (Baddeley, 2000; Chai et al., 2018). Findings on the relationship between the two domains range from moderate to strong correlations, with discrepancies likely reflecting methodological differences such as task design (Byrne et al., 2024; Kane et al., 2007), together suggesting that verbal and visuospatial WM are partially independent but related systems. For instance, verbal rehearsal strategies can counteract the rapid decay of information held in WM, supporting retention - even in visual WM tasks when items can be easily vocalized (Morey & Cowan, 2004). In children, however, visual WM may ‘lead’ this developmental chain (Hitch et al., 1989; Palmer, 2000), with a shift toward more verbally supported strategies emerging later. For instance, Gray and colleagues (2017) found that in young children’s WM was better described by a domain-general structure than by separate verbal and visuospatial components, suggesting that domain specialization itself may be a developmental achievement rather than an innate characteristic. Despite its central role in everyday cognition, the visuospatial WM has received less attention than the verbal domain, especially in developmental neuroimaging research (Klingberg, 2006; Pickering, 2001; Spencer, 2020).

These behavioral studies are supported by neural evidence too, suggesting that domain-specific development is achieved through profound structural and functional brain reorganization. Consistent with this domain distinction, verbal WM typically recruits left-lateralized, perisylvian (Broca’s and Wernicke’s) language regions, whereas visuospatial WM relies more heavily on right-lateralized fronto-parietal networks (Baddeley, 2000; Chai et al., 2018). At birth, cortical regions are interconnected by an excess of connections (Kolk & Rakic, 2022), which activity-dependent pruning refines with age into a more specialized and efficient network. Longitudinal structural MRI studies show grey matter maturation - local volume decreases also described as pruning mechanisms, following a posterior to anterior trajectory, with sensory and parietal regions maturing first and prefrontal cortex continuing to develop into adolescence (Gogtay et al., 2004; Sowell et al., 2003). Still within the primary-school years, cortical thinning in the superior parietal cortex has been linked to improved WM performance in children aged 5 to 10, whereas thinning in the anterior cingulate cortex (ACC) and the right inferior frontal gyrus specifically tracks improvements in cognitive control (Kharitonova et al., 2013).

WM is supported by a distributed frontoparietal network (FPN), comprising the dorsolateral prefrontal cortex (DLPFC), parietal cortex, and ACC, that is already present in children (aged 7-9; Klingberg, 2006; Kwon et al., 2002; Owen et al., 2005; Rosenberg et al., 2020). With development, however, activation shifts from diffuse, bilateral patterns to more focused and lateralized patterns, with the right hemisphere progressively dominating visuospatial attention and WM (Klingberg, 2006; Klingberg et al., 2002; Kwon et al., 2002).

Within this network, PFC functional maturation across childhood and adolescence underlies gradual behavioral gains in WM, inhibition, and cognitive flexibility (Chai et al., 2018), while the functionally heterogeneous ACC contributes to attentional and cognitive control via its dorsal subdivision (Botvinick et al., 2004; Heilbronner & Hayden, 2016; MacDonald et al., 2000). Parietal regions, meanwhile, are active during the WM delay period (i.e, the time between stimulus offset and the memory probe onset, during which a subject must maintain information in WM), with activity in the intraparietal sulcus (IPS) and superior parietal lobule (SPL) predicting WM capacity and load (Bray et al., 2015; Todd & Marois, 2004). These regions also contribute to attentional control and maintenance across spatial, visual, and verbal WM tasks (Dijkstra et al., 2019; Dima et al., 2014; Postle & D’Esposito, 1999). While both hemispheres contribute to attentional control, the right parietal cortex exerts stronger, more widespread influence across attentional networks (Corbetta & Shulman, 2011; Driver et al., 2004). During childhood and adolescence, the IPS shows progressive functional maturation that tracks WM capacity gains (Klingberg et al., 2002; Kwon et al., 2002).

At the network level, children as young as 7-9 years already display the same large-scale networks as adults, but within-networks integration and between-networks segregation increase with age (Fair et al., 2009; Menon, 2013; Power et al., 2010), yielding progressively more differentiated systems (Chai et al., 2014; Fair et al., 2009). A broad distinction can be drawn between task-positive networks engaged by demanding tasks (FPN and salience networks, SN) and task-negative networks such as the default mode network (DMN), active when task engagement is low (Cai et al., 2021). The DMN supports internally oriented cognition via midline regions including the precuneus/posterior cingulate cortex (PCU/PCC), medial PFC, and lateral/medial parietal cortex (Andrews-Hanna et al., 2014; Raichle, 2015). These networks interact dynamically during WM. Coordinated interactions among the SN, FPN, and DMN distinguish high from low WM load, with increasing demand strengthening frontoparietal engagement, suppressing DMN activity, and changing the influence these systems exert on one another (Cai et al., 2021; Dima et al., 2014; Wager & Smith, 2003).

In adults, DMN activity typically decreases as task-positive networks become more active during demanding tasks (Buckner et al., 2008), and age-related increases in FPN activation alongside stronger DMN deactivation across childhood and adolescence parallel WM gains (Satterthwaite et al., 2013). A similar pattern is observable earlier in childhood, but DMN-task-positive segregation is less pronounced and continues to refine with age (Chai et al., 2014; Chen et al., 2022; Fair et al., 2008), with stronger DMN-FPN segregation associated with better attention and WM performance. The PCC acts as a central hub within the DMN (Greicius et al., 2009; Leech et al., 2011), and its interaction with SN hubs such as the ACC is thought to regulate switching between internally and externally oriented brain (Anticevic et al., 2012a; Di & Biswal, 2014; Goulden et al., 2014; Greicius et al., 2009). Consistent with this network-level view, Cohen & D’Esposito (2016) describe task-based reorganization of large-scale brain networks during WM. He et al. (2023) showed that while successful WM requires rapid FPN-DMN reconfiguration, this flexible coordination is not yet fully developed in children, limiting their ability to smoothly transition from low- to high-demand states. Behavioral WM gains thus parallel the progressive specialization of frontoparietal networks (Casey et al., 2005; Klingberg, 2006; Kwon et al., 2002), underscoring the value of studying WM at the level of interacting brain networks rather than isolated regions.

Studying WM at this network level, in turn, requires methods capable of capturing directed interactions between regions. Most cognitive functions, including WM, arise not from a single brain area but from coordinated activity across distributed brain regions (Lorenc & Sreenivasan, 2021; Rezayat et al., 2022). Traditional neuroimaging studies rely on canonical activation maps, which consistently identify FPN engagement during WM tasks (Owen et al., 2005) but carry no information about interregional interactions or their directionality, leaving open the network dynamics that support WM. Effective connectivity analysis precisely captures the directed, causal influence one brain region exerts over another, and how the strength and direction of these connections vary with experimental condition (Friston et al., 2011). Human studies typically estimate effective connectivity using Dynamic Causal modeling (DCM; Stephan et al., 2010), in which neuroimaging data (e.g., fMRI, MEG) are used within a Bayesian framework to infer hidden neurobiological interactions such as the task-dependent network modulation. These estimates offer insight into the neuronal mechanisms generating observed activity (Friston et al., 2013), which allow causal inference within a predefined network, such as the FPN. Using this approach, prior studies have demonstrated a load-dependent enhancement of frontoparietal coupling during a WM task (Dima et al., 2014; Heinzel et al., 2017; Jung et al., 2018). To date, however, only a handful of studies have examined FPN effective connectivity during WM in children, and, to our knowledge, none in the visuospatial domain.

While regional brain activation during WM has been widely studied, network-level dynamics thus remain underexplored, particularly during the early school years, and within the broader framework of network interactions. Unraveling the dynamics of the functional neural correlates of WM during this important age-related period may shed light on its underlying mechanisms. The present study investigated the development of visuospatial WM in primary school children (5- to 8-year-olds), combining fMRI with DCM analysis to examine network dynamics elicited by a visual WM task. We hypothesized that (i) children would recruit the canonical FPN as observed in adults; (ii) the network would shift from a more distributed to a more specialized organization with age, marked by a stronger top-down control and right-hemisphere dominance; and (iii) segregation between task-positive and task-negative networks would increase with maturation and task demand.

## Methods

The present study used baseline (T0) data from a randomized controlled trial investigating the long-term effects of intensive artistic practice on brain maturation and executive function development in primary-school children. A detailed description of the design, interventions, and assessment procedures is provided elsewhere (James et al., 2024). Participants were assessed at three time points: baseline (T0), after one year of intervention (T1), and after two years of intervention (T2, corresponding to the end of the program). The current analyses were restricted to the baseline assessment in order to characterize age-related variability in working memory performance and its neural correlates prior to any intervention-related effects. The study was approved by the Cantonal Commission for Ethics in Human Research (CCER) and registered at ClinicalTrials.gov (NCT05912270). All procedures were conducted in accordance with the Declaration of Helsinki.

### Population

At T0, a total of 120 healthy neurotypical children aged 5-8 years (69 females; mean age = 7.29 years, SD = 0.83) were enrolled in this cross-sectional study. Only right-handed children with normal or corrected-to-normal vision and hearing, sufficient French language proficiency, and no history of neurological or psychiatric disorders were included. Exclusion criteria comprised severe neurodevelopmental disorders (e.g., severe ADHD, dyslexia), inability to complete psychometric testing or MRI incompatibility (physical or psychological). Written informed consent was obtained from parents, and assent from children, in accordance with Swiss ethics standards. Further details are provided in the study protocol (James et al., 2024).

### General procedure

All children underwent multiple preparatory steps to ensure sufficient MRI data quality. The overall procedure consisted of a mock MRI session and a training session prior to scanning. During the mock session, conducted at least three days before scanning, participants were familiarized with the MRI environment, scanner sounds, and study procedures. On the day of scanning, children completed a brief review of MRI procedures, practiced the visual working memory task (see below) until achieving at least 70% accuracy, and underwent a virtual-realitv-based MRI training session (“MRI Adventure” - https://www.pandawan.ch/pandawan-therapies/pandawan-adventure) designed to promote head stability through real-time motion feedback. Following this preparation, participants completed the MRI session.

### Working Memory Task

The visual working memory task (WM task) was adapted from Vuontela and colleagues (2003) and was performed in the MRI scanner. It consisted of a visual n-back block design paradigm adapted for children. The visual stimuli were appealing monster figures presented within a six-location spatial array arranged around a central fixation cross. The task alternated between two conditions (Figure 1A): a low-load (0-back) and a high-load (2-back) condition.

**Figure 1:**
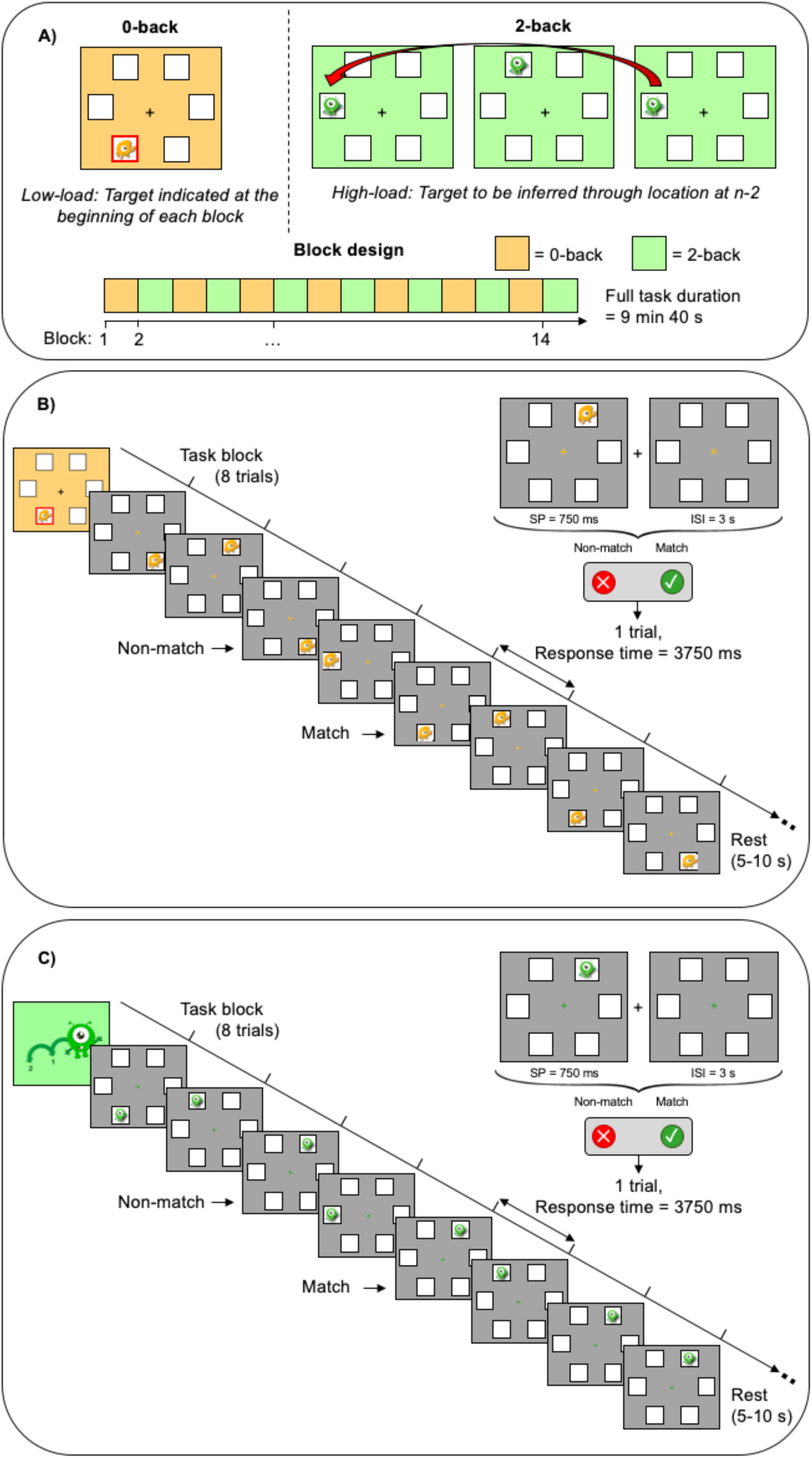
Schematic illustration of the visual working memory task. A) Top panel: visual illustration used for the instructions of the two task conditions; Bottom panel: the fixed order of the fMRI task runs. The task consisted of 14 blocks alternating between the 0-back and 2-back conditions, always starting with the 0-back condition. The 0-back condition represents the low-load condition (yellow background) and the 2-back represents the high-load condition (green background). 14 blocks alternating between 0-back and 2-back condition, starting with the 0-back condition were presented. B) Example of a 0-back block. Each screen represents one trial. A trial consists of a stimulus presentation (SP), followed by an inter-stimulus interval (ISI) (i.e. a screen only showing the fixation cross and locations). Match trials correspond to trials in which the participant can “catch” the monster (i.e., when monster is in the target location indicated at the beginning of the block), as opposed to non-match trials. C) Example of a 2-back block. Each screen represents one trial. A trial consists of a SP, followed by an ISI.. Match trials correspond to trials in which the participant can “catch” the monster (i.e., when the monster is in the same location as two trials earlier), as opposed to non-match trials. The task includes fourteen blocks, alternating between 0-back and 2-back.

At the beginning of each block, a starting slide was presented for 2000 ms to indicate the current condition (0-back or 2-back). In the 0-back condition (Figure 1B), this starting slide also indicated the block target location (one of the six locations) for 2 seconds, meaning that children must remember this target location for the entire block. On each trial, they had to determine whether the monster’s current location matched this target location shown at block onset and press the right button for a match trial, or the left button if it appeared elsewhere (i.e. non-match trial). In the 2-back condition (Figure 1C), participants were asked to press the right button if the monster appeared in the same location as two trials earlier, requiring continuous maintenance and updating of spatial information in working memory. If the location did not match the stimulus presented two trials back, they pressed the left button. Stimulus presentation (SP) lasted 750 ms, followed by a 3-second response window constituting the inter-stimulus interval (ISI). Stimuli appeared pseudo-randomly at one of the six spatial locations on a grey background screen. The task was administered in a block design that alternated cognitive load across separate 0-back and 2-back blocks to examine differences in working memory demands. There were seven blocks per condition, each consisting of eight trials, resulting in a total of 112 trials. Both conditions contained 25% match and 75% non-match trials. Blocks were separated by a rest period ranging from 5.5 to 10 seconds. The task was programmed and presented using PsychoPy 2023.2.3. (Peirce, 2007).

### Behavioral data analysis

All behavioral analyses were conducted using R software (R Core Team, 2025; RRID:SCR_001905). Performance was first assessed using d-prime (d’: signal detection theory; Green & Swets, 1966), which was computed from the hit rate (i.e. proportion of match trials correctly identified as match) and the false alarm rate (FA; i.e., the proportion of non-match trials incorrectly identified as match) both adjusted using a log-linear correction (Hautus, 1995) to correct for null or perfect performance. Overall d’ as well as d’ for separate load conditions was then calculated as the difference between the z-scored hit and FA rate. To assess whether d’ varied with age and two load condition, a linear mixed-effects model (LMM) with age, condition, and their interaction as fixed effects, and a random intercept for participant, was applied to subject-level data.

To further characterize behavioral performance at a finer grain, accuracy and RT were analyzed at the trial level. Performances were quantified for each participant by calculating accuracy (i.e. proportion of correct trials) and mean reaction time (RT) for correct trials. While the accuracy analysis included all trials, the RT analysis excluded incorrect trials and outliers (RT > 2 standard deviations from each subject’s mean; i.e. 6% of correct trials). Both measures were computed separately for the low (0-back) and high (2-back) load conditions.

A generalized linear mixed-effects model (GLMM) was applied to trial-level accuracy data, and a linear mixed-effects model (LMM) was applied to trial-level RT data, which were log-transformed to correct for their right-skewed distribution (lme4 package; Bates et al., 2015). Accuracy or RTs were modelled as dependent variables, with ‘participant’, ‘task block’, and ‘trial number’ tested as candidate random intercepts. Random effects were sequentially added to a null model to evaluate different combinations and nested structures (see Table S2). In a second stage, fixed effects and their interaction were introduced. Likelihood-ratio tests were used to compare models and select the best-fitting structure, balancing maximal predictive accuracy with minimal complexity by pruning non-significant predictors. We used ANOVA (Type III as we expected an interaction between age and condition) to assess the significance of fixed effects in the final models. Finally, to decompose the interaction effects, we estimated marginal trends using the emmeans package (Lenth et al., 2026). Specifically, we compared the estimated age slope for each condition to test the difference of the slope across conditions, for d’, accuracy and RT.

### MRI data analysis

#### MRI Data Acquisition

MRI scans were acquired on a 3-Tesla MRI scanner (Siemens Magnetom Prisma Fit, Siemens, Erlangen, Germany) using a 64-channel head/neck coil. For functional MRI acquisition during the WM task, a multiband gradient-echo EPI sequence was used (TR = 1000 ms, TE = 30 ms, flip angle = 58°, FOV = 210 × 210 × 127.5 mm, matrix = 84 × 84, 51 axial slices, slice thickness = 2.5 mm isotropic voxels, multiband/SMS factor = 3, in-plane GRAPPA factor = 2, phase partial Fourier = 7/8, bandwidth = 2290 Hz/px, echo spacing = 0.52 ms, A>>P phase encoding, number of volumes acquired = 580, total duration of 9:40 min). In addition, an MPRAGE T1-weighted scan was acquired for anatomy and normalization (TR = 2000 ms, TE = 2.49 ms, TI = 900 ms, non-selective inversion, flip angle = 9°, FOV = 230 × 230 × 166.4 mm, matrix = 288 × 288, 208 sagittal slices/partitions, GRAPPA = 2, 0.8 mm isotropic voxels).

#### MRI Preprocessing

All preprocessing used fMRIPrep 24.1.1 (Esteban et al., 2019; RRID:SCR_016216), with default, standards-compliant workflows.

##### Anatomical

Two T1-weighted (T1w) images per child were available in the dataset. Each T1w was corrected for intensity non-uniformity with N4BiasFieldCorrection (Tustison et al., 2010), as distributed in ANTs 2.5.3 (Avants et al., 2009; RRID:SCR_004757). A T1w subject template was then computed by robust within-subject registration of the T1w-corrected images using FreeSurfer mri_robust_template (7.3.2; Reuter et al., 2012), and this template served as the T1w reference. Skull stripping was performed with the Nipype implementation of antsBrainExtraction.sh (ANTs), using OASIS30ANTs as the target template. Tissue segmentation into cerebrospinal fluid, white-matter and gray-matter was performed on the brain-extracted T1w with FSL FAST (6.0.7; (Zhang et al., 2001; RRID:SCR_002823). For spatial normalization, the T1w reference was nonlinearly registered with ANTs antsRegistration to MNIPediatricAsym (TemplateFlow ID: MNIPediatricAsym, cohort-2, ages 4.5 to 8.5 years).

##### Functional

For WM task BOLD run, fMRIPrep first generated a BOLD reference for motion correction. Head-motion parameters (6 rigid-body parameters and affine transforms) were estimated with FSL MCFLIRT (Jenkinson et al., 2002) prior to any spatiotemporal filtering. The BOLD reference was co-registered to the T1w reference using FreeSurfer mri_coreg followed by FSL FLIRT with boundary-based registration cost-function (Greve & Fischl, 2009), configured with six degrees of freedom. Several confounding time-series were calculated based on the preprocessed BOLD: framewise displacement (FD), derivative of root mean square variance over voxels (DVARS), and three region-wise global signals. FD was computed using two formulations: the Power definition (absolute sum of relative motion parameters; (Power et al., 2014) and the Jenkinson definition (relative root-mean-square displacement between affines; Jenkinson et al., 2002). Both FD and DVARS were calculated using their respective Nipype implementations. The three region-wise global signals correspond to the mean BOLD signal averaged across all voxels within the CSF, WM, and whole-brain tissue masks, respectively. Confound time-series derived from head-motion estimates and global signals were expanded by adding temporal derivatives and quadratic terms for each (Satterthwaite et al., 2013). Frames exceeding a threshold of 2.0 mm FD or 6.0 standardized DVARS were flagged as motion outliers, and pooled into a single binary nuisance regressor (coded 1 for any flagged volume, 0 otherwise) rather than generated as one indicator regressor per outlier frame.

#### Univariate fMRI task-based activation analysis

Following pre-processing with fMRIPrep, a conventional general linear model (GLM) was constructed for each subject using SPM12 (*SPM12 Software - Statistical Parametric Mapping*; Penny et al., 2011; RRID: SCR_007037). Task regressors corresponding to the low-load and high-load conditions were modelled using the condition-specific onset times. These onset times were convolved with the canonical hemodynamic response function (HRF) to estimate the task-related BOLD response. To control for confounding effects, 15 nuisance regressors were included: (i) the six realignment parameters (three translations, three rotations) and their first temporal derivatives; (ii) mean white matter and cerebrospinal fluid signals, and (iii) a motion outlier regressor modeled as a binary vector to account for high-motion volumes within the GLM. Additionally, a high-pass filter (cutoff 128 s, ∼0.008 Hz) was applied to remove low-frequency drifts. The contrasts high-load > low-load and its reverse were then computed. These subject-level contrast images were subsequently entered into second-level one-sample t-tests (at the group level) analysis, to assess significant activation across participants. The resulting statistical parametric maps were thresholded using a family-wise error (FWE) correction at p < 0.05 with no cluster-extent threshold (k = 0 voxels) and overlaid on the MNI152 template for visualization.

#### Dynamic Causal Modelling (DCM) analysis

Traditional univariate activation analyses identify which brain regions are engaged by a task but do not provide information about the directional influences between regions. Dynamic causal modeling (DCM) addresses this limitation by estimating effective connectivity, that is, the causal influence that one neuronal system may exert over another.

We used DCM for fMRI (Friston et al., 2003; Zeidman et al., 2019) to model interactions within the WM network using a biophysical generative model that explains the BOLD signal via (i) hidden neuronal dynamics - bilinear state equations with intrinsic (A), modulatory (B), and input (C) couplings driven by experimental inputs (U), (ii) a hemodynamic state-space linking neural activity to BOLD via the extended Balloon-Windkessel model (Buxton et al., 1998; Friston et al., 2003), and (iii) an observation model capturing residual/noise. Parameters were estimated by variational Bayesian inversion, so ‘causality’ refers to the directed influences indicated by the fitted dynamical system Causality (Friston, 2009). To estimate these influences, we specified a model space, comprising a set of competing hypotheses about network architecture, using predefined regions of interest as nodes.

##### Selection of Regions of Interest (ROIs)

Eight ROIs were selected based on previous adult and developmental literature (Jung et al., 2018; Owen et al., 2005; Rosenberg et al., 2020) and on activation peaks identified in univariate fMRI task-based contrasts (high-load > low-load and low-load > high-load, (see 3.2.1. Task-evoked activation during the WM task). The ROIs included the bilateral middle frontal (r/lMFG), superior parietal lobules (r/lSPL), anterior cingulate cortex (r/lACC) and bilateral precuneus (r/lPCU) (Figure 2A). ROIs were defined in two steps to account for individual anatomical variability while anchoring each region to the group-level activation peak. First, a 12-mm radius sphere was centered on the group-level peak coordinate for each region (Table 1). Second, within this sphere, each subject’s local maximum was identified and used as the center of a subject-specific 6-mm radius spherical ROI. ROI timeseries were then extracted from the first-level GLM analysis: for each participant, we computed the first principal component (eigenvariate) across voxels within each ROI, resulting in a total of 960 timeseries across all participants and regions.

**Figure 2:**
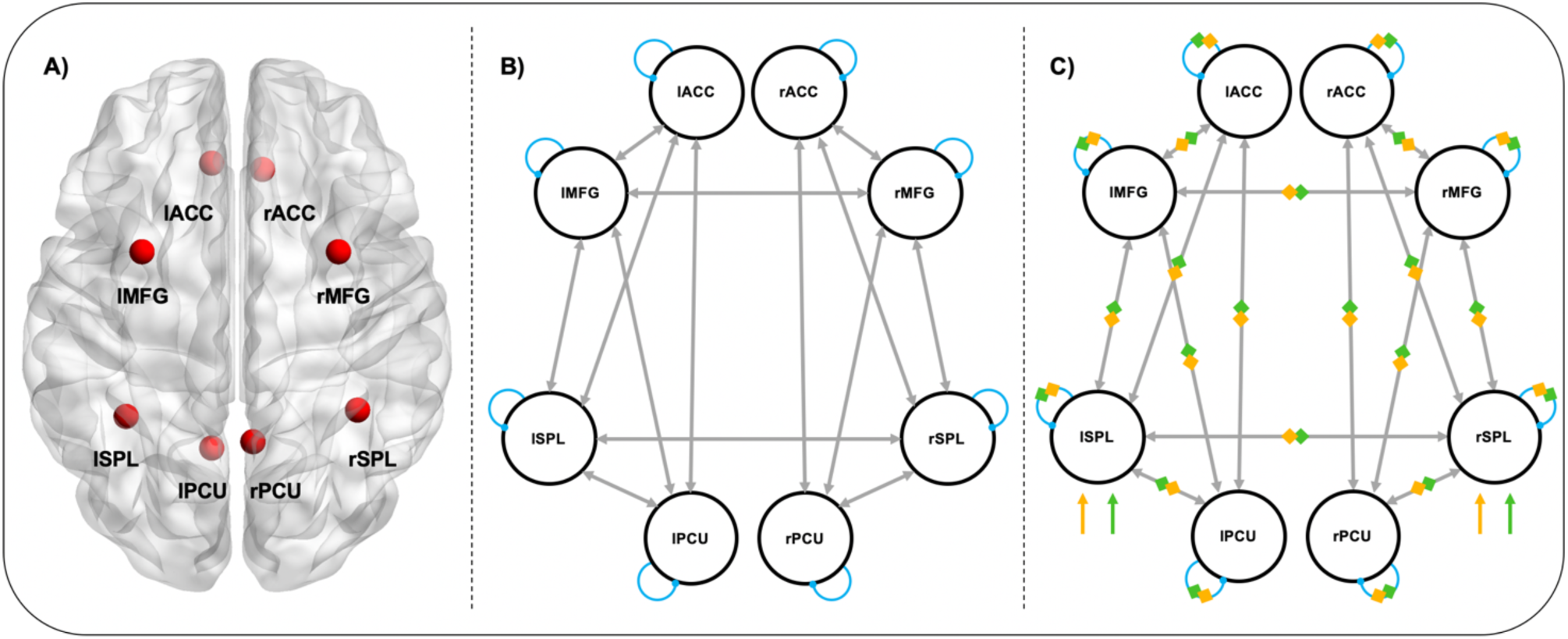
Model space for DCM analyzes on the WM task. A) ROIs used in the model, Figure created with BrainNet Viewer (Xia, 2017); B) Baseline connections (i.e. matrix A), grey arrows denote feedforward and feedback connections within each hemisphere, as well as lateral connections between homologous regions (MFG and SPL). The model also includes inhibitory self-connections (blue arrows); C) Task-dependent modulation (i.e. matrix B). The low-load (yellow diamonds) and high-load (green diamonds) conditions modulate connections between regions and inhibitory self-connections defined in matrix A. Driving inputs (i.e. matrix C) enter the model through the right and left SPL (yellow and green arrows). Abbreviations: l = left; r = right; MFG = Middle frontal Gyrus; SPL = Superior parietal sulcus; ACC = Anterior cingulate cortex and PCU = Precuneus.

**Table 1:** Sphere center of the regions of interest used in the DCM analysis. – From the contrasts of high-load > low-load condition and low > high-load condition. One-sample t-tests were performed to compare activation between the two load conditions (see 3.2.1. Task-evoked activation during the WM task). For all clusters p < 0.05 FWE. X, Y, and Z = MNI standard space coordinates (mm). Abbreviations: l = left; r = right; MFG = Middle frontal Gyrus; SPL = Superior parietal sulcus; ACC = Anterior cingulate cortex and PCU = Precuneus.

| **Label** | **X** | **Y** | **Z** |
| --- | --- | --- | --- |
| **rMFG** | 27 | 8 | 50 |
| **lMFG** | -28 | 6 | 56 |
| **rSPL** | 37 | -42 | 43 |
| **lSPL** | -36 | -44 | 40 |
| **rACC** | 7 | 34 | -7 |
| **lACC** | -10 | 34 | -10 |
| **rPCU** | 10 | -53 | 11 |
| **lPCU** | -8 | -56 | 13 |

##### Model specification

Here, we defined our full model space according to a priori hypotheses about working memory networks (Cai et al., 2021; Dima et al., 2014; Flores-Gallegos et al., 2024; Ma et al., 2012). The A-matrix represents an intrinsic (context-independent) connectivity between all ROIs (Figure 2B), modeled as full within-hemisphere coupling plus interhemispheric (‘lateral’) coupling restricted to homologous pairs only (e.g., bidirectional lSPL-rSPL), with no cross-areal interhemispheric links (e.g., bidirectional lSPL-rMFG), as callosal projections predominantly connect homologous cortical regions (Hofer & Frahm, 2006; Jarbo & Verstynen, 2015). Midline homologous connections (i.e., bidirectional rACC-lACC and rPCU-lPCU) were excluded due to their spatial proximity, potentially confounding effective estimates. Self-connections are included (on the diagonal) and log-scaled to ensure inhibitory dynamics, whereas between-region connections (off-diagonal) are expressed in Hz. All parameters were initialized with prior means of zero. The B-matrix captures task-dependent modulation of connectivity relative to matrix A. B-modulation was applied to all within-hemisphere and interhemispheric connections (including self-connections), except for midline homologous connections. This enabled us to model backward (top-down, higher order regions projecting to lower-order regions) and forward (bottom-up, lower order regions projecting to higher order) processes. In Figure 2**Erreur** **! Source du renvoi introuvable.**C, feedforward and feedback modulations are combined within a single arrow to represent bidirectional connections for visualization purposes. Lastly, the C-matrix describes task inputs to the system. Based on previous literature (Jung et al., 2018), visual input was modelled as entering bilaterally through the SPLs, consistent with the visual nature of the task (Figure 2C). Inputs were non-centered to allow the A-matrix to reflect mean connectivity.

##### First-Level Model Estimation

DCMs were first estimated for each participant, yielding subject-specific posterior estimates (i.e., model parameter estimates given the observed data) of intrinsic (A), modulatory (B), and input (C) parameters, as well as individual hemodynamic parameters (*θ*ℎ). These first-level estimates were then carried forward to the group level.

##### Group-Level Modelling

We used Parametric Empirical Bayes (PEB) to estimate group-level effects on connectivity parameters within a hierarchical Bayesian framework (Friston et al., 2016). PEB allows inter-individual variability to be decomposed into group-level effects and random effects, optimizing statistical power by pooling information across participants.

The design matrix (X) in PEB encodes subject-level covariates, with one column per predictor. All regressors were mean-centered, so that the intercept captured group mean connectivity (commonality), while the covariate columns captured between-subject effects. In our model, predictors included age and mean task accuracy. Age and accuracy were entered simultaneously so that each covariate’s effect on connectivity could be estimated adjusted for the other. Age and accuracy were only modestly correlated in our sample (Pearson r = 0.267, p = 0.003), a magnitude unlikely to introduce substantial collinearity between the two predictors and supporting their joint inclusion as a way to isolate maturational effects from performance-related effects on connectivity. This design enabled inference on (i) the group mean connectivity and its posterior probabilities and (ii) the main effects of age and accuracy on connectivity parameters under PEB (A and B matrices; Zeidman et al., 2019).

##### Model Comparison and Parameter Inference

Following model estimation, Bayesian Model Reduction (BMR) was applied to efficiently explore a nested model space by deactivating uninformative parameters. This procedure involves searching through different combinations of model parameters nested in the full model space. Reduced models were compared based on their free energy and pruned to maximize model evidence while minimizing model complexity. Next, Bayesian Model Averaging (BMA) was applied across the surviving models to obtain parameter estimates weighted by posterior model probability. We report parameters with posterior probability (Pp) > 0.99 (Zeidman et al., 2019). Although our focus was on task-induced modulations (B-matrix), we also report results for the intrinsic connectivity (A-matrix), as it represents the baseline (context-independent) network architecture upon which modulations are computed.

## Results

### Behavioral Results

The overall d’ as well as d’ for separate load conditions was computed (Table 2). We first evaluated the effects of age and load (low-load vs. high-load) condition on d’ with a LMM applied at the subject level. The model revealed a significant effect of age (*χ*²(1) = 3.874, p = 0.049) and a marginal effect of load condition (*χ*²(1) = 3.437; p = 0.064) and a significant age × condition interaction (*χ*²(1) = 3.947, p = 0.047). A post-hoc analysis of estimated marginal trends was performed to explore condition-specific slopes, revealing a positive association between d’ and age in the high-load condition (β = 0.516, SE = 0.127, 95% CI [0.266, 0.766]) though this was not significant in the low-load condition (β = 0.248, SE = 0.127, 95% CI [-0.025, 0.498]). When comparing the two slopes (low-load vs. high-load) directly, we found no significant difference (Δβ = −0.269, SE = 0.136, t= 1.97, p = 0.051).

**Table 2:**
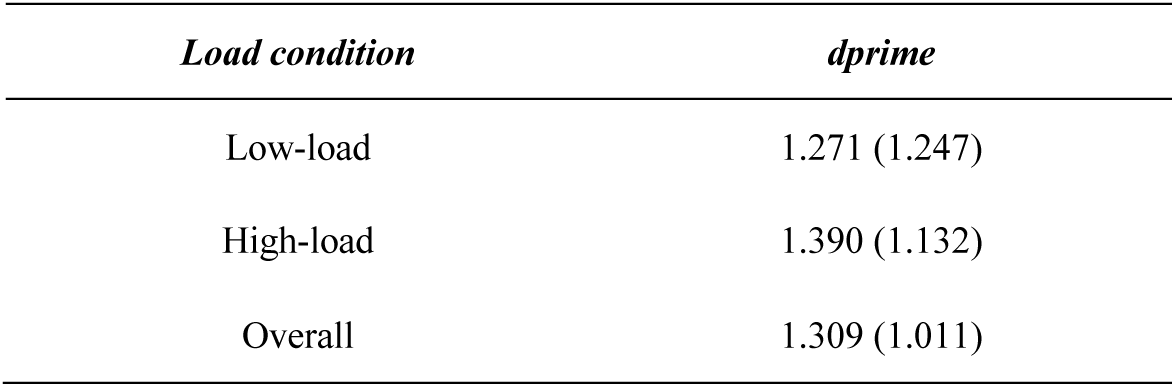
Descriptive statistics for d’ on the WM task - Mean (SD) for the low-load, high-load, and overall conditions.

| *Load condition* | *dprime* |
| --- | --- |
| Low-load | 1.271 (1.247) |
| High-load | 1.390 (1.132) |
| Overall | 1.309 (1.011) |

Then, to evaluate how accuracy and reaction times (RTs) of the WM task were affected by age and task demands, we applied a GLMM for accuracy and a LMM for RTs, at the trial level.

For accuracy, the best-fitting model, identified through likelihood-ratio comparisons, included participant, block and trial number as independent random effects, accounting for repeated measures and within-subject variability (Table S1). Fixed effects of the model included age, load condition and their interaction. The result demonstrated a significant main effect of age (*χ*²(1) = 4.773, p = 0.029), load condition (*χ*²(1) =20.07, p = 7.467e-06), and a significant age × condition interaction (*χ*²(1) = 20.57, p = 5.750e-06).

As illustrated in Figure 3A, the GLMM revealed that accuracy was positively associated with age, and that the association was more pronounced in the high-load compared to the low-load condition. Post-hoc analysis of estimated marginal trends revealed a consistent association between accuracy and age in both the low-load (β = 0.213, SE = 0.098, 95% CI [0.022, 0.404]) and the high-load conditions (β = 0.462, SE = 0.098, 95% CI [0.270, 0.654]), confirming the steeper age-accuracy association in the high-load condition (Δβ = −0.249, SE = 0.055, z= 4.54, p = 5.750185e-06).

**Figure 3:**
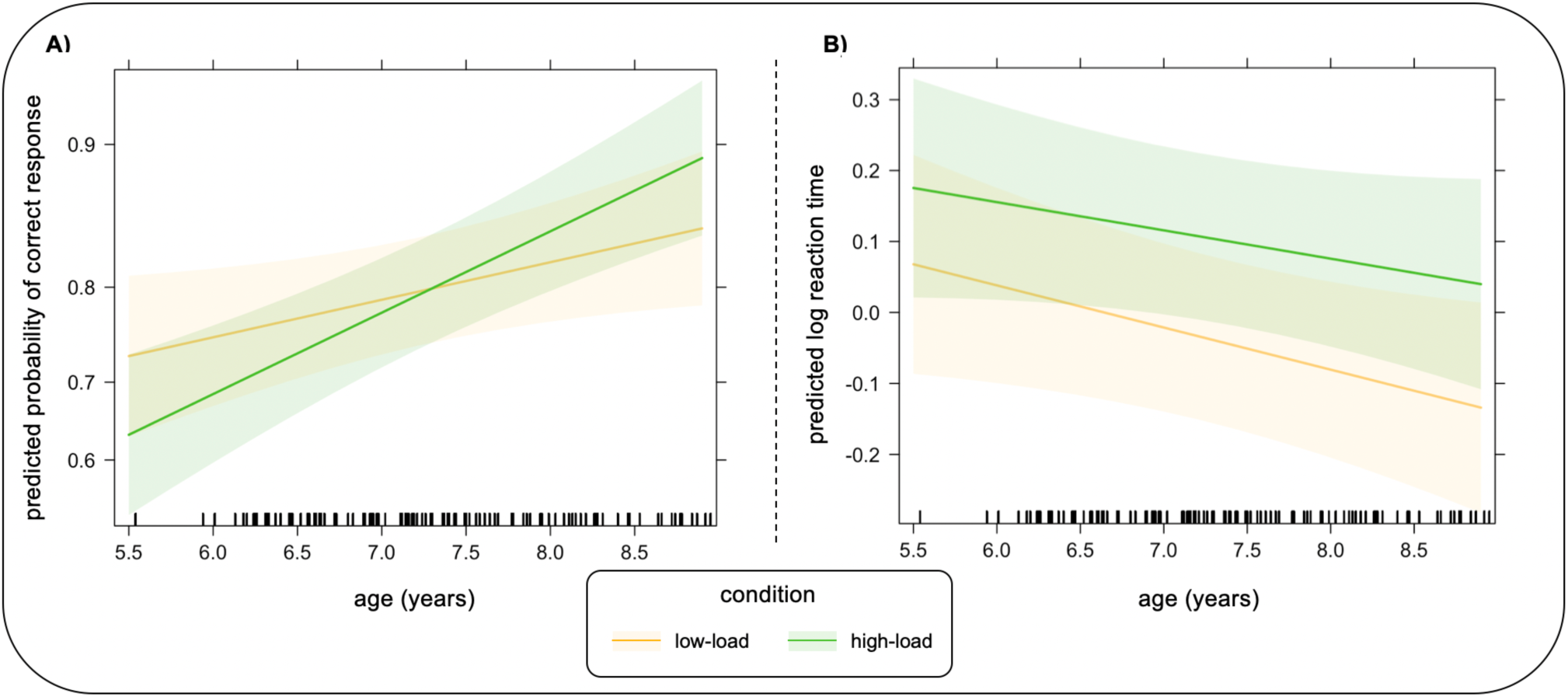
Predicted accuracy and reaction time as a function of age in the WM task. Lines represent model-estimated effects of age for each load condition (yellow = low-load, green = high-load). Shaded areas indicate 95% confidence intervals around the model predictions. (A) Predicted accuracy from a GLMM. The model revealed a significant age × condition interaction with a steeper age-related improvement under high cognitive load compared to low-load. (B) Predicted reaction time from an LMM. The model revealed significant age × condition interaction: the slope of age-related improvement (i.e. shorter reaction times) was steeper under low cognitive load compared to the high-load condition.

For RTs, the LMM included the same random and fixed effects as for the accuracy analysis (**Erreur ! Source du renvoi introuvable.**2). We found a significant main effect of age (*χ*²(1) = 4.299, p = 0.038) and a significant age x condition interaction (*χ*²(1) = 4.852, p = 0.028), indicating a stronger negative association between age and RTs in the low-load compared to the high-load condition (Figure 3B). However, the main effect of condition was not significant.

Post-hoc analysis of estimated marginal trends revealed that RT was negatively associated with age in the low-load condition (β = −0.059, SE = 0.029, 95% CI [-0.116, −0.003]) but not significantly so in the high-load condition (β = −0.04, SE = 0.029, 95% CI [-0.096, 0.016]). This negative age-RT association was significantly steeper in the low-load than in the high-load condition (Δβ = −0.02, SE = 0.009, z=-2.203, p = 0.028).

Together, these results indicate that WM performance was associated with age in different ways depending on the task demands: RTs showed a steeper negative association with age under low-load conditions, consistent with age differences in processing speed and efficiency, whereas accuracy showed a steeper positive association with age under high-load compared to low-load conditions, consistent with greater age differences in WM performance under more demanding cognitive conditions.

### FMRI results

#### Task-evoked activation during the WM task

The whole-brain second-level analysis of the high-load versus low-load (i.e. 2-back > 0-back) contrast revealed robust activation in the canonical frontoparietal control regions, including bilateral superior parietal lobule (SPL), bilateral middle frontal gyrus (MFG) and the anterior cingulate cortex (ACC) (Figure 4, hot colors). These regions are consistent with the frontoparietal network typically engaged by visual working-memory demands (Klingberg et al., 2002; Postle & D’Esposito, 1999; Zanto et al., 2011). The reverse contrast (i.e. low-load versus high-load) showed enhanced signal in regions commonly associated with lower task demand, including also the ACC, as well as the precuneus (PCU), the posterior cingulate cortex (PCC), and the bilateral hippocampi (Figure 4, cool colors). The cluster details are described in Table 3.

**Figure 4:**
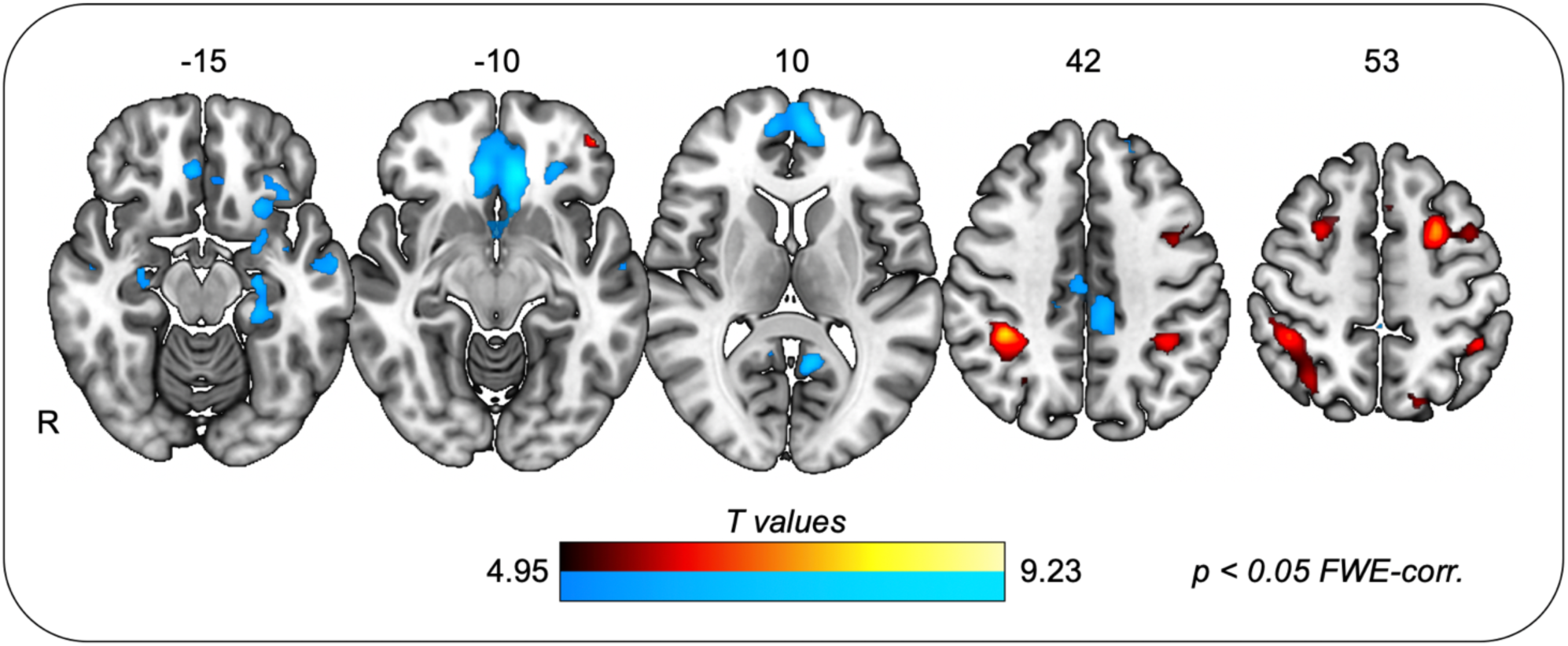
Group-level activation map for the WM task. Activation map of contrast ‘high’ load > ‘low’ load (p < 0.05 FWE-corrected). Statistical parametric maps from second-level one-sample t-tests. Hot colors indicate higher activation during high-load relative to low-load; cool colors indicate the reverse contrast. Numbers above brain slices indicate the axial slice coordinate in mm of each presented image. FWE = family-wise error.

**Table 3:** Significant activation clusters for WM contrasts. – Whole-brain contrasts high-load > low-load and low-load > high-load thresholded at p < 0.05 FWE-corrected. For each significant cluster, the table reports the peak t-value, MNI peak coordinates (x, y, z; mm), cluster size (N of voxels), and the approximate anatomical label (Neuromorphometrics atlas). Only clusters with k ≥ 100 voxels are listed (cluster extent threshold). Abbreviations: l = left; r = right; MFG = Middle frontal Gyrus; SPL = Superior parietal lobule; ACC = Anterior cingulate cortex; PCU = Precuneus; PCC = Posterior cingulate cortex; SMG = Supramarginal gyrus; HCG = Hippocampal gyrus; MTG = Middle Temporal Gyrus; FWE = family-wise error; df = degrees of freedom.

| Region | Cluster $p$-value | $N$ of voxels | $t$-value ($df = 119$) | $x$ | $y$ | $z$ |
| --- | --- | --- | --- | --- | --- | --- |
| **High > low-load contrast** |  |  |  |  |  |  |
| rSPL | 0 | 689 | 8.21 | 37 | -42 | 43 |
| lMFG | 0 | 306 | 7.95 | -28 | 6 | 56 |
| lSMG | 0 | 173 | 6.88 | -36 | -44 | 40 |
| rMFG | 0 | 123 | 6.79 | 27 | 8 | 50 |
| **Low > high-load contrast** |  |  |  |  |  |  |
| lACC | 0 | 1954 | 9.23 | -10 | 34 | -10 |
| lPCU | 0 | 165 | 7.83 | -8 | -56 | 13 |
| lPCC | 0 | 342 | 7.67 | -6 | -36 | 46 |
| lHCG | 0 | 155 | 7.57 | -26 | -16 | -20 |
| lMTG | 0 | 147 | 7.25 | -63 | -6 | -17 |

#### Effective connectivity of the WM network

Our aim was to investigate age-related differences in effective connectivity between brain regions implicated in the WM network using DCM with PEB. Briefly, we fitted a DCM model comprising eight ROIs (bilateral MFG, SPL, ACC and PCU) with bidirectional intra-hemispheric connections and interhemispheric links restricted to homologous pairs for each participant (see Figure 2A for architecture). After estimating the full models, we applied BMR to prune parameters that did not increase model evidence (i.e. free energy) by iteratively evaluating nested reductions. We then derived final parameter estimates using BMA, in which coupling parameters were averaged across the surviving reduced models and weighted by their posterior model probabilities.

Figure 5 shows the resulting networks with estimated connection strengths after thresholding at posterior probability (Pp) > 0.99. In network figures, arrow color scales with coupling strength (in Hz), with red arrows denoting excitatory and blue arrows inhibitory connections. Importantly, in DCM, between-region (extrinsic) coupling is interpreted via the polarity of the parameter; positive values signify an excitatory influence from the source to the target region while negative values represent inhibitory influence. Within-region (intrinsic) self-connections are treated differently: to ensure dynamical stability, self-connections are modeled as log-precisions with a default inhibitory prior of −0.5Hz. Therefore, the estimated values represent a deviation from this baseline inhibition. Positive self-connection values denote an increase in self-inhibition (i.e. reduced excitability) whereas negative self-connections represent a decrease in self-inhibition (i.e. disinhibition), rendering the region more sensitive to inputs and more easily recruited within the network. Mean coupling strengths for all modeled connections with Pp > 0 are summarized in Supplementary Table S2.

**Figure 5:**
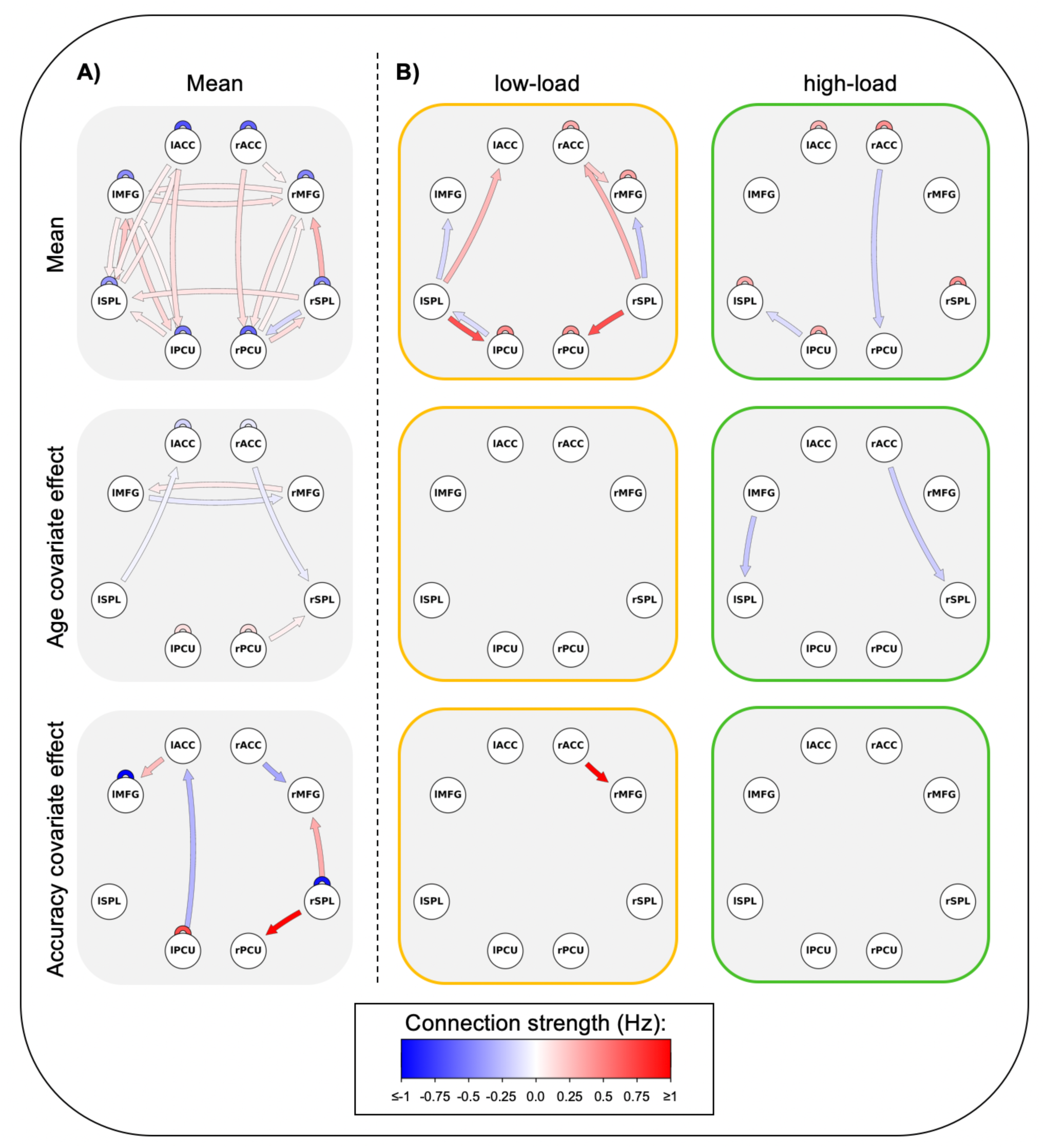
Effective connectivity of the WM network (DCM-PEB results) The model space consists of 8 nodes and two experimental task conditions (low- and high-load conditions). Edges are shown only when the posterior probability Pp > 0.99. Arrows denote direction: red = positive (excitatory) coupling, blue = negative (inhibitory) coupling. The color intensity scale represents the absolute coupling strength (units: Hz). Self-connections are plotted as deviations from the default DCM self-inhibition prior. (A) Intrinsic connectivity (A-matrix). Top: average connectivity (across both task conditions); arrows depict group-mean connections. Bottom: group-level (covariate) effects (age, accuracy) expressed as deviations from the average connectivity for the corresponding connections. (B) Task-dependent modulations (B-matrix). Yellow frame / left panels: low-load network; green frame / right panels: high-load network. Panels from top to bottom: mean condition effect, age effect and accuracy effect. Abbreviations: l = left; r = right; MFG = Middle frontal Gyrus; SPL = Superior parietal sulcus; ACC = Anterior cingulate cortex and PCU = Precuneus.

##### 1. Average effective connectivity (A-matrix)

The average effective connectivity (i.e. mean state across experimental conditions, Figure 5A), consists of a network with most coupling parameters conserved across participants (posterior probability > 0.99), most of them excitatory. We identified a fully connected architecture, with the strongest effects involving projections from SPLs to MFGs (left = 0.21 Hz; right = 0.29 Hz). Additionally, excitatory connections involving the PCUs were observed: from lMFG to lPCU (0.15 Hz), rACC to rPCU (0.13 Hz), lACC to lPCU (0.12 Hz), and rPCU to rSPL (0.13 Hz), highlighting substantial interactions with posterior midline regions. Interhemispheric connectivity was also present between homologous frontal regions (lMFG to rMFG: 0.11 Hz; rMFG to lMFG: 0.09 Hz) and between parietal regions (rSPL to lSPL: 0.10 Hz). A single negative extrinsic connection was identified from the rPCU to rSPL (−0.15 Hz). Finally, all self-connections (diagonal elements) were negative, reflecting inhibitory intrinsic dynamics. The strongest self-inhibition was observed in ACCs (left = −0.68 Hz; right = −0.62 Hz)

##### 2. Age-related modulation of effective connectivity (A-matrix covariate)

Because age and accuracy were entered simultaneously in the PEB model, the reported age effects reflect connectivity associated with age after adjusting for accuracy, and the reported accuracy effects reflect connectivity associated with accuracy after adjusting for age. Age-related effects on baseline connectivity were relatively sparse, and involved frontal interhemispheric coupling, cingulo-parietal pathways, and precuneus dynamics. Opposite age-related modulations were observed between homologous frontal regions, with a positive effect from rMFG to lMFG (0.08 Hz) and a negative effect from lMFG to rMFG (−0.08 Hz), indicating asymmetric age-related changes in interhemispheric frontal connectivity. Negative age effects were also identified between SPLs and ACCs (left = −0.04 Hz; right =−0.07 Hz), suggesting reduced cingulo-parietal interactions with increasing age. An additional positive effect was observed for the extrinsic connection from rPCU to rSPL (0.06 Hz). Finally, age modulated several self-connections: increasing excitability of ACCs (left = −0.17 Hz; right =−0.09 Hz) while reducing it in PCUs (left = 0.12 Hz; right =0.13 Hz).

##### 3. Accuracy-related modulation of effective connectivity (A-matrix covariate)

Accuracy-related effects on baseline connectivity showed up with larger values than the effects of age. The strongest effects were observed on self-connections, including negative modulations (i.e. reduced self-inhibition) of lMFG (−1.09 Hz) and rSPL (−0.92 Hz), as well as a positive modulation (i.e. increased self-inhibition) of the lPCU self-connection (0.71 Hz). Accuracy additionally modulated several extrinsic connections. Positive effects were observed from rSPL to rMFG (0.33 Hz), rSPL to rPCU (0.98 Hz), and lACC to lMFG (0.26 Hz), indicating stronger coupling associated with higher performance. In contrast, negative effects were identified for rACC to rMFG (−0.35 Hz) and lPCU to lACC (−0.29 Hz), suggesting reduced effective connectivity in these pathways with increasing accuracy.

##### 4. Task-dependent modulation of effective connectivity

Task condition (i.e. low-load versus high-load) had a significant influence on both extrinsic and intrinsic connections (Figure 5A). These parameters represent a task-dependent deviation from the average connectivity network. If a modulation (B-matrix parameter) has the same polarity as the intrinsic connection (A-matrix), the modulation enhances the existing relationship (e.g., making an excitatory connection more excitatory). If the modulation (B) has the opposite polarity to the intrinsic connection (A), it results in *down-regulation*. Specifically, if an initially excitatory connection (A > 0) is modulated by a negative task effect, the outcome depends on their relative magnitudes. If |B| > A, the connection is *suppressed,* switching the net coupling from excitatory to inhibitory.

For the low-load condition, the strongest positive modulations involved connections from the parietal to posterior midline structures (SPLs to PCUs: right = 0.69 Hz; left =0.68 Hz). Additional positive effects were observed from SPLs to ACCs (right = 0.29 Hz; left = 0.28 Hz), and from rACC to rMFG (0.21 Hz), indicating enhanced connectivity between parietal, cingulate, and frontal regions. In contrast, negative modulatory effects were observed for the parieto-frontal pathways (SPLs to MFGs: right = −0.23 Hz; left = −0.15 Hz), as well as for the lPCU to lSPL connection (−0.14 Hz). Several self-connections were also modulated by the low-load condition: positive effects were identified for rMFG (0.35 Hz), rACC (0.33 Hz), rPCU (0.43 Hz), and lPCU (0.47 Hz), indicating increased self-inhibition within frontal, cingulate, and precuneus regions.

Compared with the low-load condition, the high-load condition was characterized by a more restricted pattern of modulation, primarily involving self-connections (Figure 5A). Positive modulatory effects were observed for the self-connections of rSPL (0.43 Hz), lSPL (0.33 Hz), rACC (0.43 Hz), lACC (0.30 Hz), and lPCU (0.33 Hz), pointing to a widespread increase in the dynamics of self-inhibition during higher cognitive effort. Only two extrinsic connections were significantly modulated during high-load processing. Negative effects were observed in connections from rACC to rPCU (−0.19 Hz) and from lPCU to lSPL (−0.14 Hz), indicating reduced effective connectivity between these regions.

Overall, whereas the low-load condition induced widespread modulation of extrinsic connections linking parietal, frontal, cingulate, and precuneus regions, the high-load condition predominantly modulated intrinsic self-connections, with comparatively limited effects on inter-regional coupling.

Only one accuracy-related effect was identified during the low-load condition: a positive modulation of the connection from rACC to rMFG (1.07 Hz). In contrast, age-related effects were found only under the high-load condition. Specifically, negative age effects were observed for the connections from lMFG to lSPL (−0.22 Hz) and from rACC to rSPL (−0.20 Hz), indicating reduced effective connectivity in fronto-parietal and cingulo-parietal pathways. No age-related effects were observed during the low-load condition, and no accuracy-related effects were identified during the high-load condition. These findings indicate that age and performance selectively modulated task-dependent connectivity, with accuracy influencing cingulo-frontal interactions under low cognitive demand and age influencing fronto-parietal and cingulo-parietal interactions under high cognitive demand.

## Discussion

The present study examined how age and task performance shape effective connectivity during a visuospatial WM task in early childhood. Overall, the results confirm that children recruited a canonical, adult-like frontoparietal network; show a partial shift toward a more specialized, though not uniformly right-lateralized, architecture with age; and confirm increasing segregation between task-positive and task-negative systems with age and better task performance. Importantly, while age-related segregation of large-scale functional networks has previously been demonstrated using resting-state functional connectivity, our findings extend this developmental principle to the directed architecture of effective connectivity during active visuospatial working memory in primary school-aged children, linking greater network segregation not only to age but also to task performance.

Behaviorally, accuracy and reaction time improved with age, but the two measures diverged according to cognitive load: accuracy gains with age were steepest under high-load, whereas reaction-time gains were steepest under low-load. This load-dependent pattern was echoed in the d’ index, which showed significant effects of age and of the age × condition interaction, alongside a marginal effect of load condition alone. This dissociation is consistent with developmental accounts in which WM capacity, rather than raw processing speed, is the demand-sensitive index of maturation in middle childhood (Gathercole et al., 2004). In the low-load condition, task demands likely fall within the expected capacity across our age range, so capacity has little room to improve, making processing speed the more sensitive marker. In the high-load, task demands may exceed younger children’s capacity limits, so precision rather than speed tracks age-related change.

This pattern parallels findings from large pediatric and adolescent samples performing comparable n-back paradigms, where WM performance under increasing load has been shown to depend on the progressive engagement of executive (frontoparietal) regions together with deactivation of the DMN, and where this process tracks task performance more closely than chronological age per se (Satterthwaite et al., 2013). This behavioral dissociation allows us to draw a natural link to the effective connectivity results: if performance (not just age) indicates network maturity, then performance-related effects on connectivity would be expected to be at least as pronounced as age-related ones. This is indeed what we observe in the DCM results (see below).

At the level of univariate activation, the high- versus low-load contrast reproduced the canonical FPN signature reported in prior developmental and adult WM literature, with robust engagement of bilateral SPL, MFG, and ACC (Klingberg et al., 2002; Owen et al., 2005). The reverse contrast reproduced the well-established adult pattern of load-dependent DMN engagement (Anticevic et al., 2012a; McKiernan et al., 2003): low-load was associated with relatively higher signal in precuneus, posterior cingulate cortex, and bilateral hippocampi. These regions are typically associated with the DMN and episodic memory processing (Cavanna & Trimble, 2006; Raichle, 2015). This is not necessarily incidental to the working memory task: WM models since Baddeley (2000) posit an episodic buffer that binds temporarily maintained material with long-term/episodic memory, and the hippocampus is thought to support this kind of relational binding across a range of short delay tasks, not only long-term memory (Olsen et al., 2012). Additionally, its engagement during WM maintenance has been shown to scale with stimulus novelty (Ranganath & D’Esposito, 2001). This binding process may be more visible under low-load conditions, when executive resources are not maximally taxed, than under high-load, when executive maintenance processes dominate the BOLD signal. An alternative, non-exclusive possibility is that part of this signal reflects encoding and retrieval of the block-specific target location that must be held throughout each 0-back block, rather than (or in addition to) a load-dependent episodic-buffer process. Considered alongside the connectivity findings below, the residual DMN/hippocampal engagement observed here may nonetheless partly reflect an aspect of network organization that is still maturing at this age: the broadly defined segregation between task-positive and task-negative systems. Indeed, anticorrelation between FPN and DMN networks is thought to strengthen progressively across childhood and adolescence, with more negative (anticorrelated) FP-DMN coupling at older ages and, independently, associated with better intellectual functioning, while younger children show comparatively weaker, sometimes positive, FP-DMN coupling (DeSerisy et al., 2021). This interpretation dovetails with our DCM results, which similarly point to age-related change in average (intrinsic) fronto-parietal-precuneus connectivity, rather than in its task-dependent modulation, as the locus of age-related change.

Turning to the DCM-PEB results, the average connectivity network was dominated by strong, consistent SPL to MFG drive, alongside pronounced self-inhibition in bilateral ACC. This pattern is compatible with a bottom-up account of FPN function during visuospatial WM, in which parietal attentional/sensory regions provide the principal driving input to frontal control regions, consistent with the architecture proposed in prior DCM studies of WM networks that informed our model space (Dima et al., 2014; Ma et al., 2012). The strong ACC self-inhibition may also be consistent with this region’s proposed role as a threshold ‘gatekeeper’ that integrates conflict and performance-monitoring signals before engaging downstream control: only sufficiently strong or sustained inputs should overcome this baseline inhibition and propagate further into the network (Botvinick et al., 2004; Shenhav et al., 2013).

Age-related effects on this baseline architecture were comparatively sparse. This offers partial support for our second hypothesis; however, the pattern does not straightforwardly match the rightward lateralization reported for visuospatial attention and WM maturation in prior work (Klingberg et al., 2002; Kwon et al., 2002). The opposite-signed modulation of interhemispheric frontal coupling (positive rMFG to lMFG and negative lMFG to rMFG) suggests an asymmetric reorganization of frontal interhemispheric communication across this age-related window, while the negative age effect on SPL-ACC coupling may indicate progressive decoupling between parietal and cingulate regions. Age also modulated intrinsic self-connections: bilateral ACC became more excitable (less self-inhibited) with age, whereas bilateral precuneus became less excitable (more self-inhibited), alongside a positive age effect on the rPCU to rSPL extrinsic connection. Both findings align with the broader principle that brain networks become increasingly segregated and modular with age during childhood, a process thought to support the refinement of domain-specific processing and, ultimately, improved cognitive performance (Satterthwaite et al., 2013). This reduction in cross-regional intrinsic coupling may represent the same maturational segregation process previously described at the level of resting-state functional connectivity, here, however, observed directly in the causal (directed) architecture of the network during active task performance. Notably, ACC’s increased excitability with age echoes the pattern later seen for MFG and SPL with accuracy, while the precuneus shows the opposite trend, becoming more self-inhibited with both age and accuracy. This points to a shared mechanism: the core FPN becomes more recruitable while a DMN-linked hub becomes more tightly regulated, supporting our third hypothesis. Finally, SPL and ACC are also the regions where structural maturation (cortical thinning) has been most closely linked to WM performance gains at this age (Kharitonova et al., 2013), suggesting this functional reorganization may be scaffolded by underlying structural change.

Accuracy-related effects on baseline connectivity were more pronounced than age-related effects. Children with higher accuracy showed reduced self-inhibition in lMFG and rSPL, consistent with a more readily recruitable frontoparietal core. This was accompanied by the opposite pattern in lPCU, where higher accuracy was associated with increased self-inhibition, suggesting that better performance is not simply characterized by globally heightened excitability. Instead, this points to a more differentiated profile in which task-relevant FPN nodes become more responsive while a DMN’s hub (i.e. precuneus) becomes more tightly regulated, plausibly reflecting more effective suppression of task-irrelevant processing. This pattern is well captured by the neural efficiency framework, which proposes that more proficient cognitive performance is supported not necessarily by more neural activity, but rather by more efficiently tuned and more selectively recruitable neural systems (Neubauer & Fink, 2009). Accuracy also modulated several extrinsic pathways, with a pattern that reinforces this interpretation. Coupling strengthened with higher accuracy along three connections converging on, or reinforcing, frontoparietal communication: rSPL to rMFG, rSPL to rPCU, and lACC to lMFG. Conversely, coupling weakened with higher accuracy from rACC to rMFG and from lPCU to lACC, the latter again implicating precuneus involvement in a pattern of reduced rather than enhanced influence on the network. This reduced precuneus-to-ACC influence is consistent with the proposed role of ACC-precuneus interactions in regulating the balance between internally- and externally-oriented processing (Anticevic et al., 2012b; Greicius et al., 2009), and may reflect more effective down-regulation of internally oriented processing in higher-performing children. The hemispheric asymmetry in the ACC to MFG pathway is harder to reconcile with a single interpretive account. Based on the cingulate’s proposed role in signaling the need for control (Botvinick et al., 2004), we would have expected ACC to MFG coupling to strengthen, rather than weaken, with better performance. Yet the expected strengthening was observed only for the left-hemisphere pathway, with the right-hemisphere pathway instead weakening. We do not have a principled explanation for this reversal, and flag it as an open question. These unexpected results may reflect hemispheric specialization in ACC-MFG signaling during this age-related window. They could also be a specific feature of this sample and model space that does not generalize.

The task-dependent modulation results revealed an asymmetry that, to our knowledge, has not been clearly described in the developmental DCM literature. Low-load processing was characterized by widespread modulation of extrinsic (inter-regional) connections linking parietal, frontal, cingulate, and precuneus regions, whereas high-load processing was characterized predominantly by modulation of intrinsic self-connections, with comparatively limited inter-regional change. This asymmetry is broadly consistent with accounts of coordinated, load-dependent reorganization across frontoparietal, salience, and default mode systems (Cai et al., 2021; Cohen & D’Esposito, 2016), while suggesting a developmentally specific refinement. Rather than scaling inter-regional recruitment continuously with load, as might be expected of a fully mature network, children’s networks may depend on a more segregated repertoire in which reconfiguration operates primarily at low demand and gives way to local, self-connection adjustments once demand increases. This would align with evidence that the rapid FPN-DMN reconfiguration needed to transition smoothly between low- and high-demand states is not yet fully mature in children (He et al., 2023). This is a testable hypothesis for future work directly modeling the relationship between intrinsic coupling strength and the locus of task-dependent modulation.

Finally, age and accuracy effects on task-dependent modulation showed a double dissociation by load: accuracy effects emerged only under low-load (a single rACC to rMFG modulation), whereas age effects emerged only under high-load (reduced lMFG to lSPL and rACC to rSPL coupling). This pattern complements the behavioral dissociation reported above where age-related accuracy gains were steepest under high-load, suggesting that age and individual performance capture partly distinct aspects of load-dependent network organization. Together, the behavioral and connectivity results converge on a coherent narrative. WM development in this age range may be characterized by load-dependent differences in network organization. This phenomenon may be expressed as a speed-accuracy dissociation at the behavioral level, and as a dissociation between ‘where’ in the network (self- versus inter-regional connections) and ‘under which condition’ (low- versus high-load) age and performance leave their respective signatures at the neural level.

These findings should be interpreted considering several limitations. First, although baseline data were drawn from a longitudinal cohort, the present analyses are cross-sectional, restricting our developmental claims to age-related variability rather than within-subject change. The assessment of two additional timepoints will allow these hypotheses to be tested prospectively. Second, the narrow age range studied (5-8 years) limits generalization to a window already past the earliest emergence of WM, and may underestimate age effects relative to studies spanning childhood through adolescence. Third, our focus on the visuospatial domain, while addressing a relative gap in the developmental neuroimaging literature, leaves open how these findings relate to verbal or domain-general WM processes. The latter caveat is sharpened by evidence that WM in young children may be better described by a domain-general than a domain-specific structure (Gray et al., 2017). Fourth, our DCM model space, while grounded in prior literature, necessarily reflects a priori choices about node selection and permissible connections (e.g., restricting interhemispheric links to homologous pairs, fixing driving inputs to the SPLs). Alternative model spaces, for instance, incorporating additional DMN hubs beyond the precuneus, might reveal a different balance of intrinsic and modulatory effects. Finally, while DCM affords a substantial advantage over functional connectivity in estimating directed, causal influences, parameter estimates remain dependent on the specified generative model and should be interpreted as model-conditional rather than as direct physiological measurements.

In conclusion, our findings show that visuospatial WM development in primary school-aged children is characterized not simply by stronger recruitment of frontoparietal regions, but by increasingly differentiated effective connectivity between task-positive and task-negative systems. By demonstrating that this organization varies with both age and task performance, and that its expression differs according to cognitive load, the present study extends previous developmental functional connectivity findings to the directed architecture of brain networks during active WM. Together, these results highlight effective network segregation and load-dependent reconfiguration as candidate mechanisms supporting the development of efficient working memory during childhood.

## Acknowledgements

C.E.J, D.M., M.K. and J.R. acknowledge support from the Swiss National Science Foundation (SNSF 214977). This study was conducted using the imaging platform and virtual reality platform of the Brain and Behaviour Laboratory (BBL), Cognitive and Affective Neuroimaging Section of the Center for Biomedical Imaging (CIBM MRI UNIGE).

