## Supplementary material for "Frontoparietal effective connectivity supports working memory in early childhood"

*Table S1:* ***Linear mixed models performed on the WM task: random effect model specification*** *– For each variable, we defined a model containing the most appropriate random effects. Random effects were introduced sequentially, and model fit was assessed through Likelihood Ratio Tests by comparing each model to simpler alternatives (see methods). Model notation follows Rs lme4 package convention: 1|Subject = random participant intercept; 1 | Block = random block intercept; 1 | Trial = random trial intercept; the '+' operator denotes additive random effects, whereas the '/' operator indicates nesting (i.e., lower-level units grouped within higher-level units). The final 'random effect' model is underlined. AIC: Akaike Information Criterion; LRT: Likelihood Ratio Test; Chisq: Chi-square. P-value significance codes: '***' < 0, 0.001; '**' < 0.01; '*' < 0.05; '.' < 0.1.*

| **Dependent variable** | | ***Model fit*** | | ***LRT test against simpler model*** |
|---|---|---|---|---|
| **Random effects** | ***AIC*** | ***Deviance*** | ***df*** | ***Chisq*** |
| **Accuracy** | | | | |
| 0 | - | - | - | |
| 1\|Subject | 9498.5 | 9494.5 | - | - |
| 1\|subject_id + 1\|num_block | 9476.0 | 9470.0 | 1 | 7.476e-07 *** |
| 1\|subject_id/num_block | 9452.6 | 9446.6 | 0 | 0 |
| <u>1\|subject_id + 1\|num_block + 1\|num_trial</u> | 9352.8 | 9344.8 | 1 | < 2.2e-16 *** |
| 1\|subject_id + 1\|num_block/num_trial | 8283.5 | 8275.5 | 0 | 0 |
| 1\|subject_id/num_block + 1\|num_trial | 9329.4 | 9321.4 | 0 | 0 |
| 1\|subject_id/num_block/num_trial | 9454.6 | 9446.6 | 0 | 0 |
| **Reaction time** | | | | |
| 0 | - | - | - | - |
| 1\|Subject | 12163 | - | 0 | - |
| | | 6078.4 | | |
| 1\|subject_id + 1\|num_block | 11861 | 11853 | 1 | < 2.2e-16 *** |
| 1\|subject_id/num_block | 11897 | 11889 | 0 | 0 |
| <u>1\|subject_id + 1\|num_block + 1\|num_trial</u> | 10816 | 10806 | 1 | < 2.2e-16 *** |
| 1\|subject_id + 1\|num_block/num_trial | 10826 | 10816 | 0 | 0 |

*Table S2:* ***Descriptive statistics of dynamic causal modelling connection strengths.*** *Source and Target indicate directionality of the connections. Connections with a posterior probability of 0 are marked as '–'or omitted when absent across an entire row. Abbreviations: Common. = Commonalities (i.e. value of connection strength), Pp = Posterior probability, l = left; r = right; MFG = Middle frontal Gyrus; SPL = Superior parietal sulcus; ACC = Anterior cingulate cortex and PCU = Precuneus.*

| | | ***Average connectivity*** | | ***Age covariate*** | | ***Accuracy covariate*** | |
|---|---|---|---|---|---|---|---|
| ***Source*** | ***Target*** | ***Common.*** | ***Pp*** | ***Common.*** | ***Pp*** | ***Common.*** | ***Pp*** |
| | | | | ***Matrix A*** | | | |
| rMFG | rMFG | -0.4918 | 1 | - | 0 | - | 0 |

| | | | | | | | |
|---|---|---|---|---|---|---|---|
| rMFG | lMFG | 0.0912 | 1 | 0.0800 | 1 | - | 0 |
| rMFG | rPCU | 0.0955 | 1 | - | 0 | -0.2586 | 0.609547 |
| lMFG | rMFG | 0.1122 | 1 | -0.0838 | 1 | - | 0 |
| lMFG | lMFG | -0.4526 | 1 | - | 0 | -1.0928 | 1 |
| lMFG | lSPL | 0.0557 | 1 | - | 0 | - | 0 |
| lMFG | lPCU | 0.1542 | 1 | - | 0 | - | 0 |
| rSPL | rMFG | 0.2913 | 1 | - | 0 | 0.3317 | 1 |
| rSPL | rSPL | -0.4698 | 1 | - | 0 | -0.9173 | 1 |
| rSPL | lSPL | 0.1018 | 1 | - | 0 | - | 0 |
| rSPL | rACC | - | 0 | 0.0261 | 0.622987 | - | 0 |
| rSPL | rPCU | -0.1520 | 1 | - | 0 | 0.9854 | 1 |
| lSPL | lMFG | 0.2126 | 1 | - | 0 | - | 0 |
| lSPL | rSPL | 0.0299 | 0.611883 | - | 0 | - | 0 |
| lSPL | lSPL | -0.4086 | 1 | - | 0 | -0.2243 | 0.55733 |
| lSPL | lACC | 0.0721 | 1 | -0.0440 | 1 | - | 0 |
| rACC | rMFG | 0.0529 | 1 | -0.0289 | 0.59121 | -0.3464 | 1 |
| rACC | rSPL | - | 0 | -0.0720 | 1 | - | 0 |
| rACC | rACC | -0.6157 | 1 | -0.0855 | 1 | - | 0 |
| rACC | rPCU | 0.1376 | 1 | - | 0 | - | 0 |
| lACC | lMFG | 0.0239 | 0.613033 | - | 0 | 0.2656 | 1 |
| lACC | lSPL | 0.0600 | 1 | - | 0 | 0.1352 | 0.541407 |
| lACC | lACC | -0.6826 | 1 | -0.1671 | 1 | - | 0 |
| lACC | lPCU | 0.1291 | 1 | - | 0 | - | 0 |
| rPCU | rMFG | 0.0503 | 1 | - | 0 | - | 0 |
| rPCU | rSPL | 0.1358 | 1 | 0.0578 | 1 | - | 0 |
| rPCU | rACC | - | 0 | -0.0130 | 0.474333 | - | 0 |
| rPCU | rPCU | -0.5939 | 1 | 0.1309 | 1 | - | 0 |
| lPCU | lMFG | 0.0406 | 1 | - | 0 | - | 0 |
| lPCU | lSPL | 0.0782 | 1 | - | 0 | - | 0 |
| lPCU | lACC | - | 0 | - | 0 | -0.2958 | 1 |
| lPCU | lPCU | -0.5243 | 1 | 0.1239 | 1 | 0.7080 | 1 |
| | | | ***Matrix B – low-load*** | | | | |
| rMFG | rMFG | 0.3520 | 1 | - | 0 | - | 0 |
| rSPL | rMFG | -0.2263 | 1 | - | 0 | - | 0 |
| rSPL | rACC | 0.2936 | 1 | - | 0 | - | 0 |
| rSPL | rPCU | 0.6886 | 1 | - | 0 | - | 0 |
| lSPL | lMFG | -0.1536 | 1 | - | 0 | - | 0 |
| lSPL | lSPL | 0.0992 | 0.5370 | - | 0 | - | 0 |
| lSPL | lACC | 0.2805 | 1 | - | 0 | - | 0 |
| lSPL | lPCU | 0.6844 | 1 | - | 0 | - | 0 |
| rACC | rMFG | 0.2056 | 1 | - | 0 | 1.0722 | 1 |
| rACC | rSPL | - | 0 | - | 0 | 0.8357 | 0.78995295 |

| | | | | | | | |
|---|---|---|---|---|---|---|---|
| rACC | rACC | 0.3266 | 1 | - | 0 | - | 0 |
| rACC | rPCU | 0.1802 | 0.829542649 | - | 0 | - | 0 |
| rPCU | rSPL | -0.0848 | 0.774642946 | - | 0 | - | 0 |
| rPCU | rPCU | 0.4260 | 1 | - | 0 | - | 0 |
| lPCU | lSPL | -0.1382 | 1 | - | 0 | - | 0 |
| lPCU | lPCU | 0.4675 | 1 | - | 0 | - | 0 |
| ***Matrix B – high-load*** | | | | | | | |
| lMFG | lMFG | 0.1632 | 0.76678361 | - | 0 | 1.0578 | 0.70821899 |
| lMFG | lSPL | - | 0 | -0.2225 | 1 | - | 0 |
| rSPL | rSPL | 0.4305 | 1 | - | 0 | - | 0 |
| rSPL | rACC | - | 0 | - | 0 | -0.5028 | 0.54955475 |
| lSPL | lSPL | 0.3332 | 1 | - | 0 | - | 0 |
| rACC | rSPL | -0.1155 | 0.75921121 | -0.2025 | 1 | - | 0 |
| rACC | rACC | 0.4308 | 1 | - | 0 | - | 0 |
| rACC | rPCU | -0.1909 | 1 | - | 0 | - | 0 |
| lACC | lACC | 0.2950 | 1 | - | 0 | - | 0 |
| lPCU | lSPL | -0.1354 | 1 | - | 0 | - | 0 |
| lPCU | lPCU | 0.3322 | 1 | - | 0 | - | 0 |
